# Discrete Subfields and Continuous Gradients Coexist: A Multi-Scale View of Hippocampal Organization

**DOI:** 10.1101/2025.08.19.671141

**Authors:** Nichole R. Bouffard, Morgan D. Barense, Morris Moscovitch

## Abstract

The human hippocampus is studied via two competing frameworks: one dividing it into discrete anatomical subfields with distinct computational processes, and another describing it as a continuous, functional gradients along the anterior-posterior and medial-lateral axes. How these organizational principles relate to one another, particularly regarding intrinsic neural timescales of the hippocampus, remains unknown. Here, we used high-resolution resting-state fMRI to investigate how single-voxel autocorrelation, a measure of intrinsic neural timescale, maps onto hippocampal subfields. We found evidence for a hybrid organization. First, consistent with our predictions, we observed significantly higher autocorrelation (longer timescales) in the subiculum compared to the other subfields. Contrary to our hypotheses, we found that CA1, which is implicated in integration, had low autocorrelation whereas CA2/3 and CA4DG, which are linked to pattern separation, had intermediate autocorrelation. Second, we discovered that the overarching anterior-posterior and medial-lateral gradients of autocorrelation were recapitulated *within* each individual subfield. Finally, data-driven clusters of autocorrelation values aligned more strongly with the continuous gradients than with the discrete anatomical boundaries, particularly in the right hemisphere. These results suggest that the discrete and continuous views of hippocampal organization are not mutually exclusive but coexist across different spatial scales. We therefore propose a new comprehensive model of hippocampal function that integrates both its modular, subfield-specific properties and its graded, large-scale organization.

## Introduction

Our ability to form rich, detailed memories of life events depends critically on the hippocampus. A long-standing question in neuroscience is how the intricate anatomical structure of this brain region gives rise to its complex mnemonic functions. To address this, researchers have largely adopted one of two influential frameworks. The first, a discrete approach, views the hippocampus as a collection of distinct subfields, each performing a specialized computational role. The second, a continuous approach, characterizes the hippocampus by its gradients of functional organization, primarily along the long axis. While both frameworks are supported by substantial evidence, they offer conflicting views on hippocampal organization. In this study, we aim to bridge these perspectives by investigating the intrinsic neural timescales, or autocorrelation, of individual voxels across discrete subfields and continuous anatomical axes.

Decades of anatomical research in animals and humans have identified distinct hippocampal subregions, or subfields, based on their cytoarchitectonic and receptor-architectonic features. The human hippocampus is divided into several subfields along the anterior-posterior and medial-lateral axes: CA1, CA2/3, CA4, dentate gyrus (DG), the subiculum, and stratum radiatum, lacunosum, and moleculare (SRLM) (Amunts et al., 2005; Palomero-Gallagher et al., 2020). This discrete anatomical framework has been foundational for cognitive neuroscientists, who have proposed theoretical and computational models ascribing distinct roles to these subfields. A prominent example is the Complementary Learning Systems model, which characterizes the functional roles of the subfields in relation to an entorhinal-hippocampal circuit that supports learning and memory (McClelland et al., 1995; O’Reilly et al., 2014). In this model, DG and CA3 make memories distinct from one another using orthogonal neural patterns to learn individual episodes, whereas CA1 performs statistical learning using overlapping neural patterns to represent and integrate across similar memories (Schapiro et al., 2016, 2017). While supporting evidence for this model comes from examinations of neural patterns of fMRI BOLD signal in the hippocampus, little is known about the fluctuations of signal over time in the subfields and whether they can be differentiated by their intrinsic neural timescales.

The subiculum is often excluded from the computational models of the human hippocampal subfield circuitry. Some researchers consider the subiculum to be the primary output region that relays processed signals from CA1-4/DG to other cortical regions via direct projections to the entorhinal cortex (O’Mara, 2006; Zeidman & Maguire, 2016). However, a growing body of evidence from both rodent and human studies suggest the subiculum performs computations distinct from the other subfields and has its own subregions. The most medial portion of the subiculum is further broken down into the pre- and parasubiculum (Zeidman et al., 2015). Investigations of the pre/parasubiculum in the rodent hippocampus have found grid, border, and head direction cells that are important for spatial processing (Boccara et al., 2010; Cacucci et al., 2004; Taube, 1995). Studies in humans also find that the subiculum is distinctly involved in spatial processing, and it has been proposed that the subiculum may be specialized for the holistic processing of scenes (Dalton & Maguire, 2017; Zeidman et al., 2015). Specifically, Zeidman, Lutti, et al. (2015) propose that the anterior pre/parasubiculum are particularly important for binding together elements of a scene, which contributes to the construction of an internal model or representation of scenes in memory. This integrative process might be characterized by longer neural timescales, or slower fluctuations in neural activity overtime, yet prior fMRI work in humans has been limited to examinations of univariate activity or neural patterns, leaving open the question of how neural timescales might differentiate the distinct computational processes of the subfields.

In contrast to the discrete framework of hippocampal subfields, other research has employed a continuous approach to study hippocampal function and organization. Human fMRI studies have revealed gradients of functional connectivity, particularly along the anterior-posterior axis of the hippocampus (Genon et al., 2021; Przeździk et al., 2019; Raut et al., 2020; Vos de Wael et al., 2018). The graded change along the long axis is not specific to functional connectivity measures, and a robust anterior-posterior gradient has also been found in studies of intrinsic hippocampal temporal autocorrelation (Bouffard, Golestani et al., 2023; Brunec, Bellana et al., 2018). Recently a secondary gradient of functional connectivity (Vos de Wael et al., 2018) and intrinsic temporal autocorrelation (Bouffard, Golestani et al., 2023) has been found along the medial-lateral axis of the hippocampus. Crucially, these functional gradients do not neatly map onto the anatomical boundaries of the hippocampal subfields (Genon et al., 2021). The disconnect between continuous gradients and anatomical subfields is most prominent in studies that have applied data-driven parcellation methods, which are agnostic to anatomical boundaries and instead are based on functional connectivity of the hippocampus (Barnett et al., 2019; Chase et al., 2015; Cheng et al., 2020; Dalton, McCormick, Luca, et al., 2019; Ge et al., 2019; Masouleh et al., 2020; Plachti et al., 2019; Robinson et al., 2015; Zhong, 2019). The resulting data-driven parcellations consistently are aligned along the anterior-posterior and medial-lateral axes functional axes rather than the underlying anatomical subfields. This suggests that signal dynamics are not constrained to anatomically defined subregions and instead vary throughout the hippocampus in a way that might more closely relate to the functional neural gradients (Genon et al., 2021).

These two frameworks are not necessarily mutually exclusive. Evidence from human and rodent work suggests that there are neural and molecular gradients within each individual subfield. For example, prior work from human neuroimaging found anterior-posterior gradients of functional connectivity within the subiculum, CA1-3, and CA4/DG (Vos de Wael et al., 2018). Single unit recordings in the rodent hippocampus have also revealed distinct neural gradients within subfields, defined by a dorsal-ventral gradient of signal firing patterns in CA1 and CA3 (Barnes et al., 1990; Jung et al., 1994; Keinath et al., 2014; Kjelstrup et al., 2008; Komorowski et al., 2013; Leutgeb et al., 2004; Maurer et al., 2005). Further, research investigating the gene expression of cells within the rodent hippocampus also found distinctions along the long axis, where DG and CA1 are segregated into three major molecular domains (dorsal, intermediate, ventral) and CA3 can be divided into 9 molecular domains (Strange et al., 2014). The neural and molecular gradients found within the hippocampal subfields suggest that we should not reduce examinations of hippocampal subfields down to units that serve one functional role and instead should consider that neural processes may vary along the long axis. While there have been numerous studies in rodents which have revealed gradients of neuronal signaling in hippocampal subfields, human fMRI studies of neural gradients within hippocampal subfields have been limited (but see Chase et al., 2015; Dalton, McCormick, Luca, et al., 2019; and Vos de Wael et al., 2018). An aim of the current study was to expand this literature and examine whether hippocampal gradients of intrinsic timescales, measured as single voxel autocorrelation, are recapitulated in each subfield, or whether the gradients differ among the subfields, or both.

Here we analyzed a high-resolution resting state fMRI dataset to examine the relationship between intrinsic neural timescales and the anatomical subfields of the hippocampus. We first used a discrete approach and investigated whether the subfields could be differentiated by distinct neural timescales. In accord with the prominent model of hippocampal subfields that treats subfields as homogenous units, we predicted that the neural timescales would parallel the cognitive processes attributed to each subfield. For example, pattern separation might be a process that is supported by shorter timescales, therefore we expected lower autocorrelation in DG. Generalization and integration might be supported by longer timescales, and therefore we expected higher autocorrelation in CA1. For the subiculum, it is possible that longer neural timescales and slower neural fluctuations are necessary for binding and integrating scene elements to create a holistic representation. We predicted, therefore, higher autocorrelation in the subiculum than in the other subfields. Another possibility is that gradients of autocorrelation are recapitulated within individual subfields. We tested this hypothesis by employing a continuous approach and examining gradients of autocorrelation along the anterior-posterior and medial-lateral axes of each subfield. Lastly, we applied data-driven clustering to the voxels and computed the overlap between data-driven clusters and the anatomical subfields.

## Methods

### Participants

We analyzed resting state fMRI data from 25 participants (15 female; age range: 22-36) from the Human Connectome Project (HCP) dataset. This dataset is a subset of the HCP 1200 Subjects data release, which consists of data from 1200 participants who were scanned using the full HCP imaging protocol on a 3T (structural) and a 7T (functional) scanner. All subject recruitment procedures and informed consent forms, including consent to share de-identified data, were approved by the Washington University Institutional Review Board (IRB) (Glasser et al., 2016). The present analysis of this dataset was approved by the University of Toronto research ethics board.

### Scanning parameters and preprocessing

A high-resolution 3D anatomical scan was collected on a Siemens 3T “Connectome Skyra” (Washington University in St. Louis, USA) for each subject (T1-weighted sequence, FOV 224x224 mm, flip angle = 8 degrees, voxel size of 0.7 x 0.7 x 0.7 mm). This subset of HCP subjects were additionally scanned on a Siemens Magnetom 7T MR scanner (Center for Magnetic Resonance (CMRR) at University of Minnesota in Minneapolis, MN) where high-resolution resting state fMRI was collected. Resting state data were collected using a gradient-echo EPI pulse sequence (TR = 1000 ms, TE = 22.2 ms, 85 slices with 1.6 mm thickness, FOV = 208 x 208 mm, voxel size = 1.6 mm isotropic, Flip angle = 45, Multiband factor = 5, Scan time = 16 minutes) with eyes open and relaxed fixation on a projected bright cross-hair on a dark background (and presented in a darkened room). Every participant completed four scanned runs. Oblique axial acquisitions alternated between phase encoding in a posterior-to-anterior (PA) direction in runs 1 and 3, and an anterior-to-posterior (AP) phase encoding direction in runs 2 and 4. For the present study we only analyzed the first scanned run from every participant, which was collected with a posterior-to-anterior phase encoding direction.

Initial fMRI preprocessing steps already applied to the downloaded data (labelled as “minimally preprocessed data” by HCP) included gradient distortion correction, motion correction (based on FLIRT), field map distortion correction, brain extraction, nonlinear registration to T1-weighted image and MNI standard space (1.6 mm resolution), intensity normalization, and bias field removal (Glasser et al., 2016; Smith et al., 2013; Van Essen et al., 2013). The data were further preprocessed (labeled as “clean data” by HCP) by running ICA using FSL’s MELODIC function with automatic dimensionality estimation limited to a maximum of 250. ICA is able to separate multiple signal and noise components, including head motion and physiological signals. These components are fed into the FIX in FSL (Griffanti et al., 2014; Salimi-Khorshidi et al., 2014), which classifies components into “good” vs. “bad”. Bad components are then removed from the data in a “non-aggressive” manner, in which only the unique variance associated with the bad components is removed from the data. Twenty-four confound timeseries derived from the motion estimation (the 6 rigid-body parameter timeseries, their backwards-looking temporal derivatives, plus all 12 resulting regressors squared) are also regressed out. No spatial smoothing was applied, however there is a degree of smoothing that occurs during the data interpolation which is required for registration of the fMRI data into the standard MNI space. To eliminate high frequency noise and artifacts, fMRI signals are low-pass filtered using MATLAB IIR Butterworth filter (designfilt function in Signal Processing Toolbox) with cutoff frequency of 0.1 Hz before computing the single voxel autocorrelation.

### Hippocampal subfield segmentation

Hippocampal subfield masks for each participant were created using the automated HippUnfold software and protocol (DeKraker et al., 2022). Subfield masks were created for each participant using their native space T1w anatomical scan. The HippUnfold applies a surface-based topological unfolding technique to create precise segmentations. A histologically derived unfolded reference atlas (3D BigBrain) was used to segment and label the following hippocampal subfields: subiculum, CA1, CA2, CA3, CA4, dentate gyrus (DG), and the stratum radiatum, lacunosum, and moleculare (SRLM). The resolution of functional fMRI that is typically recommended for delineation between CA2/CA3 and CA4/DG is 1.5 mm (advice from Dr. Rosanna Olsen). Here our resting state functional data was 1.6 mm, therefore we chose to combine CA2 and CA3 segments and CA4 and DG segments for the analyses in the paper. The resulting subfield segments we analyzed were: CA1, CA2CA3, CA4DG, SRLM, and subiculum.

To compute single voxel autocorrelation, the native space anatomical masks (voxel size 0.7 mm isotropic) needed to be warped to the resting state functional scans (voxel size 1.6 mm isotropic). Minimally processed and cleaned functional scans were co-registered to MNI space using FNRIT (nonlinear). We summed all the subfield masks together to create a whole hippocampal mask. We then registered each mask (the whole hippocampus mask and each individual subfield mask) to every participant’s functional resting state scan. The whole hippocampal mask was used to compute single voxel autocorrelation throughout the entire hippocampus and the subfield masks were used to extract single voxel autocorrelation values from each subfield.

### Single voxel autocorrelation method — Computing single voxel autocorrelation

For each voxel inside of the whole hippocampal mask, unbiased autocorrelation (as described in Bouffard, Golestani et al., 2023) was calculated. Briefly, the timecourse of a single voxel’s activity was correlated with itself shifted by a temporal lag, the length of 1 TR (1000 ms). This was repeated four times, shifting the timecourse forward by 1 lag until a maximum temporal shift of 4000 miliseconds was reached. The autocorrelation (AC) computed for each lag was stored in a vector. The autocorrelation vector (single voxel autocorrelation vector) contained 4 values (one single voxel autocorrelation for each lag). This approach resulted in a single voxel autocorrelation vector for each voxel. We used the hippocampal subfield masks derived from each subject’s native space to extract groups of voxels and computed the average single voxel autocorrelation for each subfield. We computed the average single voxel autocorrelation by only taking the autocorrelation value at lag1 for each voxel (following the methods Bouffard, Golestani et al., 2023). We re-ran analyses by examining all four lags separately, as well as by averaging all four lags. These analyses resulted in similar results to our lag1-only analysis, therefore we only report the results for the lag1 analysis. Autocorrelation vectors (all four values) were used to compute autocorrelation clusters.

### Visualizing single voxel autocorrelation gradients in subfields

We examined gradients of single voxel autocorrelation along the anterior-posterior and medial-lateral axes of each subfield. To visualize autocorrelation gradients, we sliced the hippocampus along the anterior-posterior axis with 1.6 mm slices and averaged the autocorrelation (lag1 only) across voxels on each slice. We did this at the individual level, for each subfield separately. We then averaged across individuals and plotted the group-level values to visualize the gradient. Because we used participant-specific subfield segmentations, there were anatomical differences among participants and some participants had larger/longer hippocampi, which resulted in some participants with more slices along the anterior-posterior axis. To be able to make comparisons across participants, we included only anterior-posterior slices which had data from 75% or more of participants (18 participants). We repeated these steps in the medial-lateral direction by taking 1.6 mm slices in along the medial-lateral axis (from left to right) and plotting the average autocorrelation per slice for each subfield. We included only medial-lateral slices that had data from 75% or more of participants.

### Computing individual-level autocorrelation clusters

To compute the data-driven autocorrelation clusters at the individual-level, we followed the method from Bouffard, Golestani et al., 2023. Briefly, we used participant-specific whole hippocampal masks to extract the autocorrelation vectors. The Euclidean distance between the single voxel autocorrelation vectors of each voxel pair in the mask was calculated to generate a similarity matrix for the whole mask. This similarity matrix was used to generate hippocampal clusters using the modularity optimization algorithm proposed by (Blondel et al., 2008; Wickramaarachchi et al., 2014), which does not require a preset number of clusters and instead estimates the optimum number of clusters from data. An autocorrelation cluster map for left and right hippocampus was created for each participant.

### Computing overlap between subfields and autocorrelation clusters

To evaluate the overlap between subfields and autocorrelation clusters, we computed the proportion of voxels that overlapped between each subfield every autocorrelation cluster. We computed the overlap between the anatomical subfield segmentations that had been registered to each participant’s resting state scan (voxel size 1.6 mm isotropic) and each participant’s autocorrelation cluster map derived from the resting state scan (voxel size 1.6 mm isotropic) (Figure 3, right). For a given cluster, we summed the number of voxels belonging to a specific subfield and then divided by the total number of voxels in the cluster to get the proportion of overlap between that subfield and the cluster. We repeated this process for each subfield separately. The proportion of overlapping voxels between subfields and autocorrelation clusters was calculated within each individual and each hemisphere separately then averaged to get the group-level proportion of overlap.

### Computing group-level autocorrelation clusters

We computed group-level autocorrelation clusters. Visually examining autocorrelation clusters at the individual-level is noisy. It is easier to evaluate the predominant spatial distribution of clusters by examining the group-level maps, which are less noisy. Group-level autocorrelation maps allow us to label the clusters according to their anatomical location (e.g., anterior-medial cluster, posterior-lateral cluster). To compute the group-level autocorrelation cluster maps, we first co-registered individual resting state scans to 2 mm MNI standard space, which required resampling of the resting state scans from 1.6 mm to 2 mm. We recalculated the single voxel autocorrelation for each participant using the resampled resting state scans using the bilateral 2mm MNI hippocampal mask from the Brainnetome atlas (Fan et al., 2016). We applied data-driven clustering to the single voxel autocorrelation vectors at the individual-level to create individual autocorrelation cluster maps. We then averaged the similarity matrices of all participants to create the group-level autocorrelation cluster map.

## Results

### Average single voxel autocorrelation within subfields

Our first analysis approach was a discrete approach, and we examined each subfield as an individual unit. We averaged the autocorrelation across all voxels within each hippocampal subfield (lag1 only). We ran a linear mixed effects model on the average autocorrelation with subfield (subiculum, CA1, CA2CA3, CA4DG, SRLM) and hemisphere (left, right) as predictors. We included participant as the random intercept in the random effects term. We found a main effect of subfield (F(4, 216) = 38.36, p <0.001) and a main effect of hemisphere (F(1, 216) = 82.34, p <0.001) (Figure 1). There were no significant interactions. For the post hoc analysis of the main effects, we used the Kenward-Roger method to calculate the degrees of freedom and the Tukey method to correct for multiple comparisons. Analysis of the main effect of subfield revealed that autocorrelation in the subiculum was greater than that of the other subfields (CA1 (t(216) = 11.40, p < 0.001), CA2CA3 (t(216) = 5.56, p < 0.001), CA4DG (t(216) = 8.38, p < 0.001), and SRLM (t(216) = 9.09, p < 0.001)). We also found that CA1 had lower autocorrelation than the subiculum (t(216) = 11.40, p < 0.001), CA2CA3 (t(216) = 5.84, p < 0.001), and CA4DG (t(216) = 3.02, p < 0.05). CA1 was not significantly lower than the SRLM. Analysis of the main effect of hemisphere revealed that across subfields, the right hemisphere had greater autocorrelation than the left hemisphere (t(216) = 9.07, p < 0.001).

**Figure 1.**
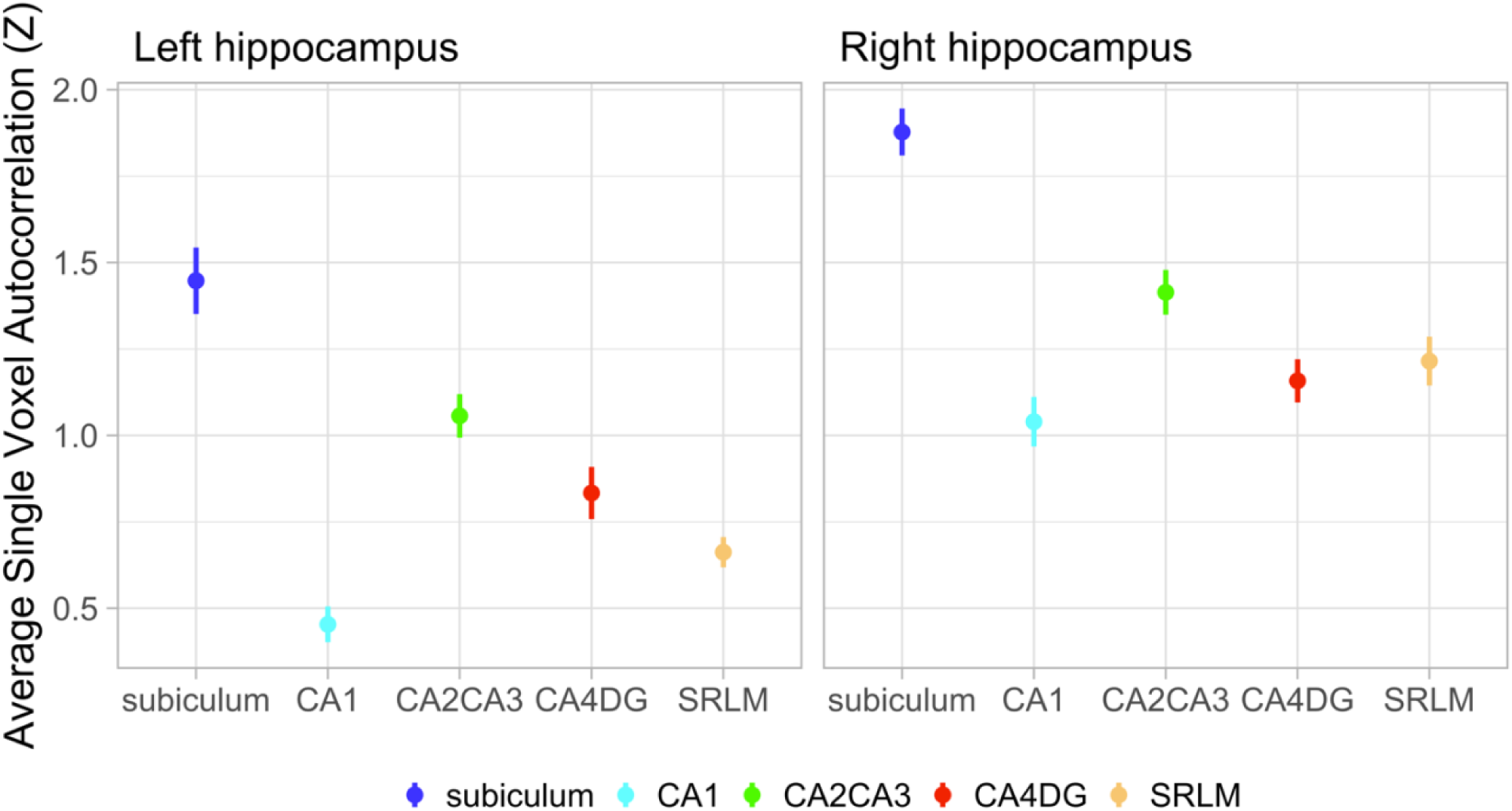
Average single voxel autocorrelation in hippocampal subfields. Average single voxel autocorrelation was computed for each subfield and then Z-scored within each participant. Plot depicts the average autocorrelation for each subfield averaged across all participants. Error bars denote standard error.

### Single voxel autocorrelation gradients in subfields

Our second analysis approach was a continuous approach, and we examined gradients of single voxel autocorrelation along the anterior-posterior and medial-lateral axes of each subfield. We found that in the right hemisphere, there was an anterior-posterior gradient in all subfields, where low autocorrelation values were in the posterior hippocampus and autocorrelation increased as slices progressed more anteriorly (Figure 2). We also found that the subiculum had the highest autocorrelation compared to other subfields along the long axis. There was a slight decrease in autocorrelation in the most anterior slices of all the subfields except CA4DG. This is likely due the decrease in number of voxels per slice in the most anterior slices in conjunction with there being lower autocorrelation in the lateral hippocampus, which could drive down the overall average autocorrelation on slices that have fewer voxels. In the left hemisphere we did not find the expected anterior-posterior gradient. Across all subfields in the left hemisphere the autocorrelation had minimal change in value along the long axis. There was also a decrease of autocorrelation in the most anterior slices.

**Figure 2.**
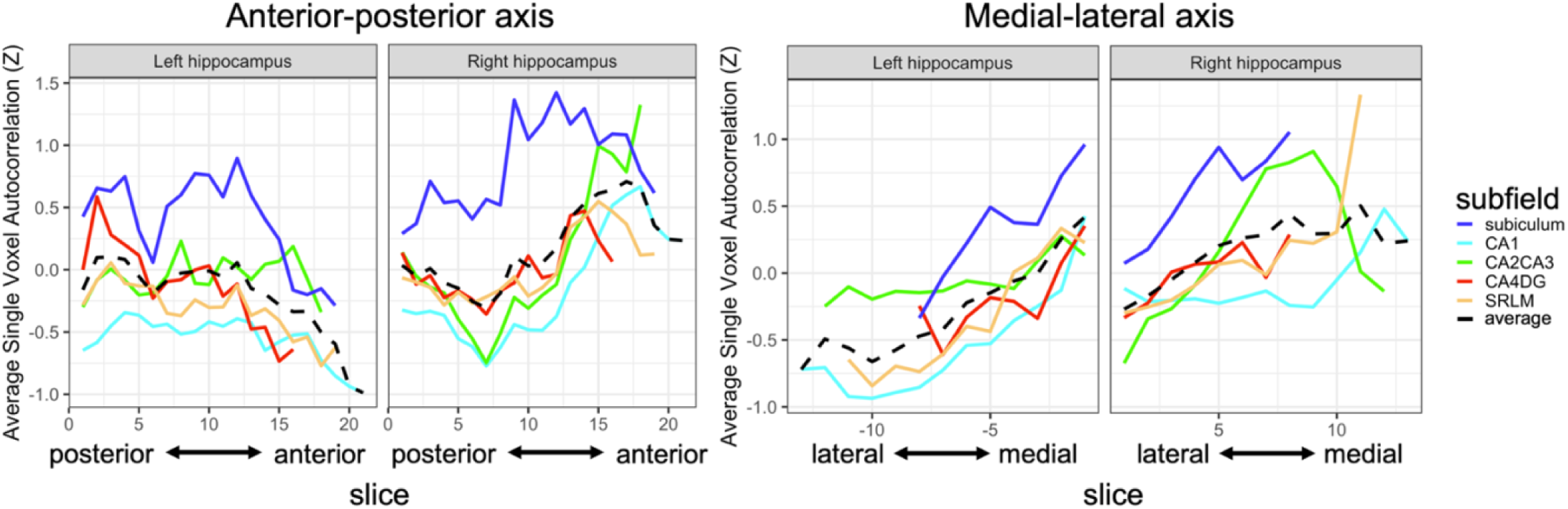
Single voxel autocorrelation gradients along the anterior-posterior and medial-lateral axes. X axis is the average autocorrelation across all voxels on each hippocampal slice. Y axis represents slices in the anterior-posterior (left) and medial-lateral (right) axes. Colored lines represent the group average autocorrelation across participants for each subfield. Black dashed line is the average of autocorrelation across all subfields.

In both hemispheres, there was a medial-lateral gradient in all subfields, where low autocorrelation values were in the lateral hippocampus and increased as slices moved more medially (Figure 2). Here we were able to replicate our past work which found a high-to-low gradient of autocorrelation along the medial-lateral axis of the hippocampus.

### Individual-level autocorrelation clusters

Data-driven autocorrelation clusters were computed for each participant. Every participant had three autocorrelation clusters in each hemisphere. There was one cluster with high autocorrelation, one with medium autocorrelation, and one with low autocorrelation. The anatomical location of the autocorrelation clusters can be hard to label based on visual examination of the individual-level cluster maps because they are noisier and cluster definitions are less clear compared to the group-level maps (see Figure 3 for an autocorrelation cluster map from an example participant). In our prior work (Bouffard, Golestani et al., 2023; Coughlan et al., 2023), we have previously identified the high autocorrelation cluster located in the anterior-medial hippocampus, the medium autocorrelation in the intermediate hippocampus, and the low autocorrelation cluster in posterior-lateral hippocampus. We therefore labeled the three autocorrelation clusters found in the present study as anterior-medial, intermediate, and posterior-lateral. We also computed the group-level autocorrelation clusters and evaluated the anatomical location based on the group-level maps (see *Group-level Autocorrelation Clusters* section below).

**Figure 3.**
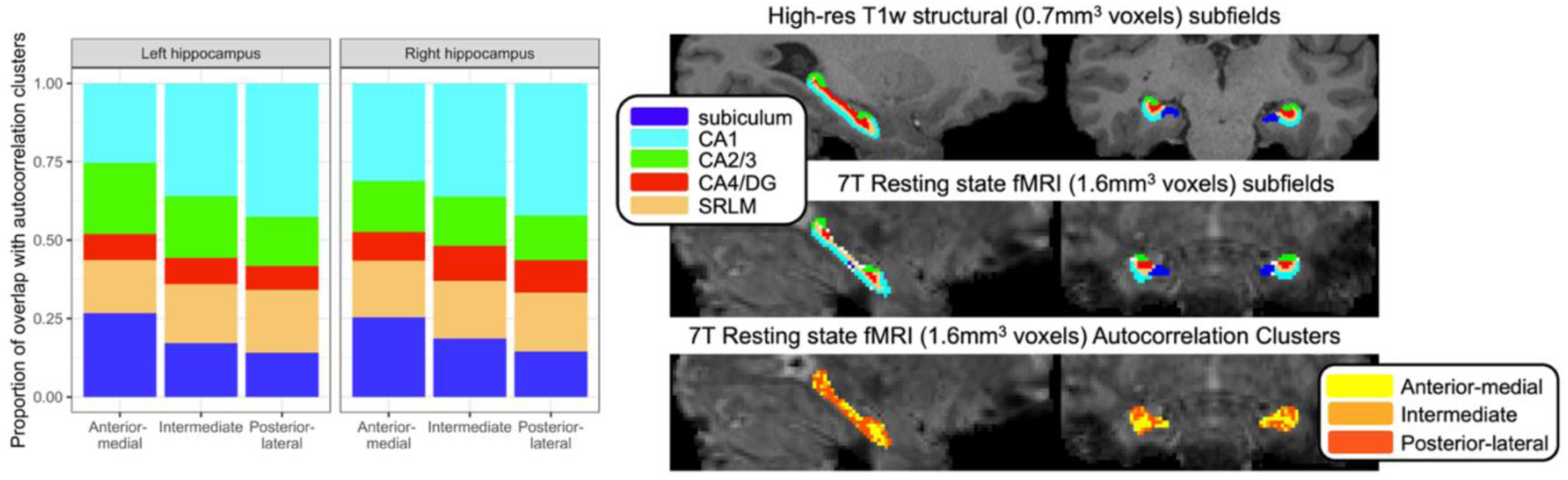
High resolution subfield masks vs. data-driven autocorrelation clusters. Left. Proportion of overlap of subfields with autocorrelation clusters. Right. Hippocampal subfield segmentation and autocorrelation clusters for an example participant.

### Overlap between subfields and autocorrelation clusters

To examine how much overlap the autocorrelation clusters had with the underlying anatomical subfields, we calculated the proportion of overlap between each subfield with the three autocorrelation clusters. Within each subfield, we compared the proportion of overlap with the anterior-medial compared to proportion of overlap with the intermediate and posterior-lateral clusters. For the following comparisons we used the Kenward-Roger method to calculate the degrees of freedom and the Tukey method to correct for multiple comparisons. We found that in both hemispheres, subiculum had greater overlap with the anterior-medial cluster compared to the intermediate (Left: t(691)= 8.69, p < 0.001; Right: t(691)= 6.39, p < 0.001) and posterior-lateral cluster (Left: t(691)= 11.42, p < 0.001; Right: t(691)= 9.97, p < 0.001) (Figure 3, left). We also found that in both hemispheres CA1 had greater overlap with the posterior-lateral cluster compared to the intermediate (Left: t(691)= 6.42, p < 0.001; Right: t(691)= 6.16, p < 0.001) and compared to the anterior-medial cluster (Left: t(691)= 16.42, p < 0.001; Right: t(691)= 10.63, p < 0.001). For the rest of the subfields (CA2CA3, CA4DG, and SRLM), there were similar proportions of voxels in the anterior-medial, intermediate, and posterior-lateral clusters (Left: ts < 3.22, ps > 0.08; Right ts < 1.79, ps > 0.89) with the exception of CA2CA3 in the left hemisphere, where there was a greater proportion of voxels in the anterior-medial cluster (t(691)= 6.22, p < 0.001) and intermediate cluster (t(691)= 3.61, p < 0.024) compared to the posterior-lateral cluster.

### Group-level autocorrelation clusters

In prior work, autocorrelation values were organized into three clusters along the anterior-posterior and medial-lateral axes of the hippocampus (Bouffard, Golestani, et al., 2023). Here we aimed to replicate these past findings by computing the group-level autocorrelation clusters in a high-resolution resting state dataset. We found that three autocorrelation clusters were produced in each hemisphere (Figure 4, top). Autocorrelation in the right hemisphere was consistent with our past work and the clusters were organized into an anterior-medial cluster (Cluster 1) of high autocorrelation values, posterior-lateral cluster of low autocorrelation values (Cluster 3), and an intermediate cluster (Cluster 2) of intermediate autocorrelation values (Figure 4). Autocorrelation in the left hemisphere was also organized into three clusters, ranging from high to low autocorrelation, however the clusters did not follow an anterior-medial to posterior-lateral organization. In the left hemisphere we found that low autocorrelation cluster (Cluster 3) was located in the anterior-lateral hippocampus instead of in the posterior-lateral hippocampus and the high autocorrelation cluster (Cluster 1) was located in both the anterior-medial and posterior-medial hippocampus instead of only the anterior-medial hippocampus. There seemed to be preservation of the medial-lateral organization, where the low autocorrelation cluster (Cluster 3) was located more laterally and the high autocorrelation cluster (Cluster 1) was located more medially, however the anterior-posterior organization was not consistent with our past work. We were only able to partially replicate our past work – the right hemisphere followed the predicted organization of clusters along the anterior-posterior and medial-lateral axes, but the left hemisphere did not, particularly along the anterior-posterior axis.

**Figure 4.**
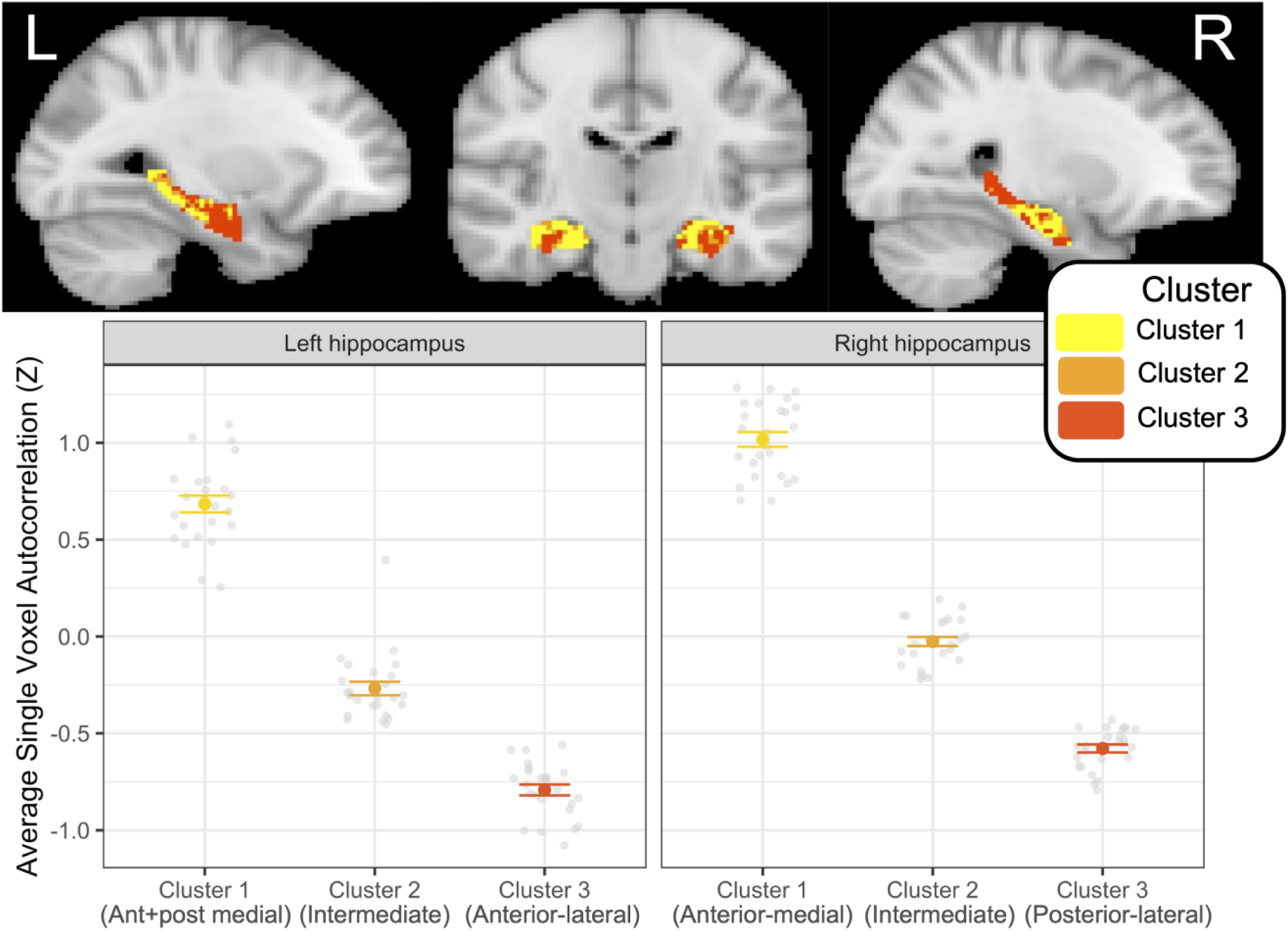
Top. Group-level Autocorrelation Clusters in MNI standard space (2 mm). Sagittal and coronal views of the group-level autocorrelation map for the left and right hemisphere. There were three group-level autocorrelation clusters (Cluster 1, Cluster 2, Cluster 3). The clusters in the right hemisphere followed the anterior-medial, intermediate, and posterior-lateral spatial organization, consistent with prior work. The left hemisphere did not have the same organization, and instead Cluster 1 was in both anterior-medial and posterior-medial hippocampus, and Cluster 3 was in the anterior-lateral hippocampus. Bottom. Average single voxel autocorrelation in group-level autocorrelation clusters. Average autocorrelation was calculated by averaging the autocorrelation values across all the voxels in each autocorrelation cluster. Average autocorrelation was then Z-scored within each participant. Grey circles represent the average autocorrelation for each participant and colored circles represent the group-average autocorrelation. Error bars denote standard error.

## Discussion

In this study we investigated the relationship between single voxel autocorrelation and anatomical hippocampal subfields. We analyzed a high-resolution 7T resting state fMRI dataset, which allowed us to examine autocorrelation of individual voxels within the smaller substructures of the hippocampus. We first took a discrete approach by examining average autocorrelation in each of the subfields. Consistent with our predictions we found that the subiculum had significantly higher autocorrelation than the other subfields. Analysis of the other subfields revealed that the autocorrelation differed across subfields in an unexpected way. CA1, which has been linked to integration processes, had low autocorrelation and CA2/3 and CA4DG, which have been linked to pattern separation processes, had intermediate autocorrelation. We investigated this further by examining the distribution of single voxel autocorrelation throughout the hippocampus and found continuous gradients of autocorrelation along the anterior-posterior and medial-lateral axis of each subfield. Given that gradients of autocorrelation are recapitulated within individual subfields, we propose that discrete models of the hippocampal subfields that assign a single functional role to each subfield might be overlooking the possibility that the functions performed by each subfield are differentiated and vary along the long axis.

In theoretical and computational models of the human hippocampus, the subfields are described as individual units performing computation processes, without any differentiation of their function along the long axis of the subfields (Dimsdale-Zucker et al., 2018; Kyle et al., 2015; Schapiro et al., 2017; Yassa & Stark, 2011). There is evidence based on functional connectivity that the anterior CA1 and subiculum can be differentiated from posterior CA1 and subiculum, and that anterior versus posterior segments of the subfields contribute to different memory networks (Libby et al., 2012; Ranganath & Ritchey, 2012; Ritchey et al., 2015). Despite evidence that there are graded changes of functional connectivity along the long axis of the hippocampus, computational models of the hippocampal subfields have not been updated to account for long axis gradients. One reason why the discrete and continuous models of the hippocampus are difficult to integrate is that there have been limited studies that examine functional distinctions along the long axis of subfields that are related to behavior, which is due to the limitations of spatial resolution in fMRI (but see Zeidman, Lutti, et al., 2015). The current study analyzed resting state data and thus cannot make direct links to behavior, but we can integrate our gradient findings with the predominant computational model of the hippocampal subfields to incorporate long axis organization and make predictions about the relationship to behavior.

In the hippocampal-entorhinal circuit, information enters from the entorhinal cortex and is sent to the DG, CA3, and CA1. In the first pathway, information is processed via the trisynaptic pathway (TSP), where the sparsely coded signal from DG and CA3 is then passed to CA1, which has overlapping neural patterns, and sent back out to the entorhinal cortex (Figure 5; Schapiro et al., 2017). In this model there are distinct neural codes in the DG and CA3 but overlapping neural codes in CA1. In the second pathway, the monosynaptic pathway (MSP), information is sent from entorhinal cortex to CA1 where there are distributed neural codes, which then project back out to the entorhinal cortex (Figure 5). It is unclear from this model how the sparse neural code from DG and CA3 is maintained after being passed through CA1 and back to the entorhinal cortex. Researchers have hypothesized that the relative influence that CA3 and the entorhinal cortex has on the neural patterns in CA1 is dynamic, and depending on the memory being retrieved the CA1 will either reflect overlapping patterns driven by stronger input from entorhinal cortex or will demonstrate more distinct sparse firing patterns based on strong input from CA3 (Guzowski et al., 2004).

**Figure 5.**
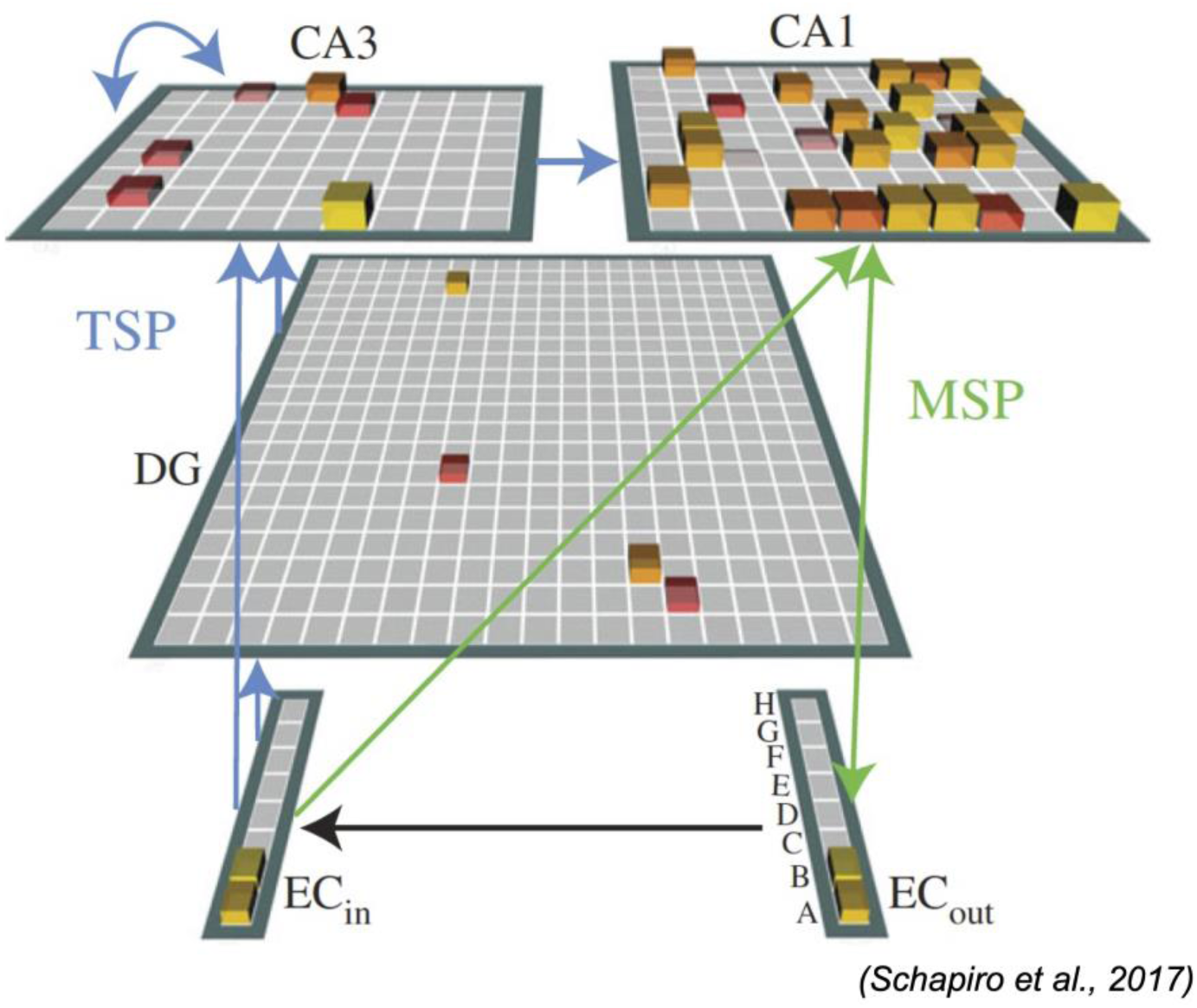
**Figure adapted from Schapiro et al. (2017)** and illustrates the computational model of the hippocampal subfields. Information enters from the entorhinal cortex and is sent to the DG, CA3, and CA1. In the first pathway, information is processed via the trisynaptic pathway (TSP), where the sparsely coded signal from DG and CA3 is then passed to CA1, which has overlapping neural patterns, and sent back out to the entorhinal cortex. In this model there are distinct neural codes in the DG and CA3 but overlapping neural codes in CA1. In the second pathway, the monosynaptic pathway (MSP), information is sent from entorhinal cortex to CA1 where there are distributed neural codes, which then project back out to the entorhinal cortex.

We build on the previous model by proposing that CA1 signal output qualitatively differs along the long axis (Figure 6). For example, high autocorrelation in the anterior CA1 supports the integrative function typically associated with CA1, whereas low autocorrelation in the intermediate and posterior CA1 supports the more sparsely coded information coming from the CA3 and DG (Schapiro et al., 2017). Under this proposal, the posterior CA1 maintains the distinct patterns sent from CA3 and DG via rapid fluctuations in neural activity, whereas anterior CA1 integrates information and with overlapping neural patterns that are supported by slow fluctuations in neural activity and passed on to the subiculum and entorhinal cortex (Figure 6).

**Figure 6.**
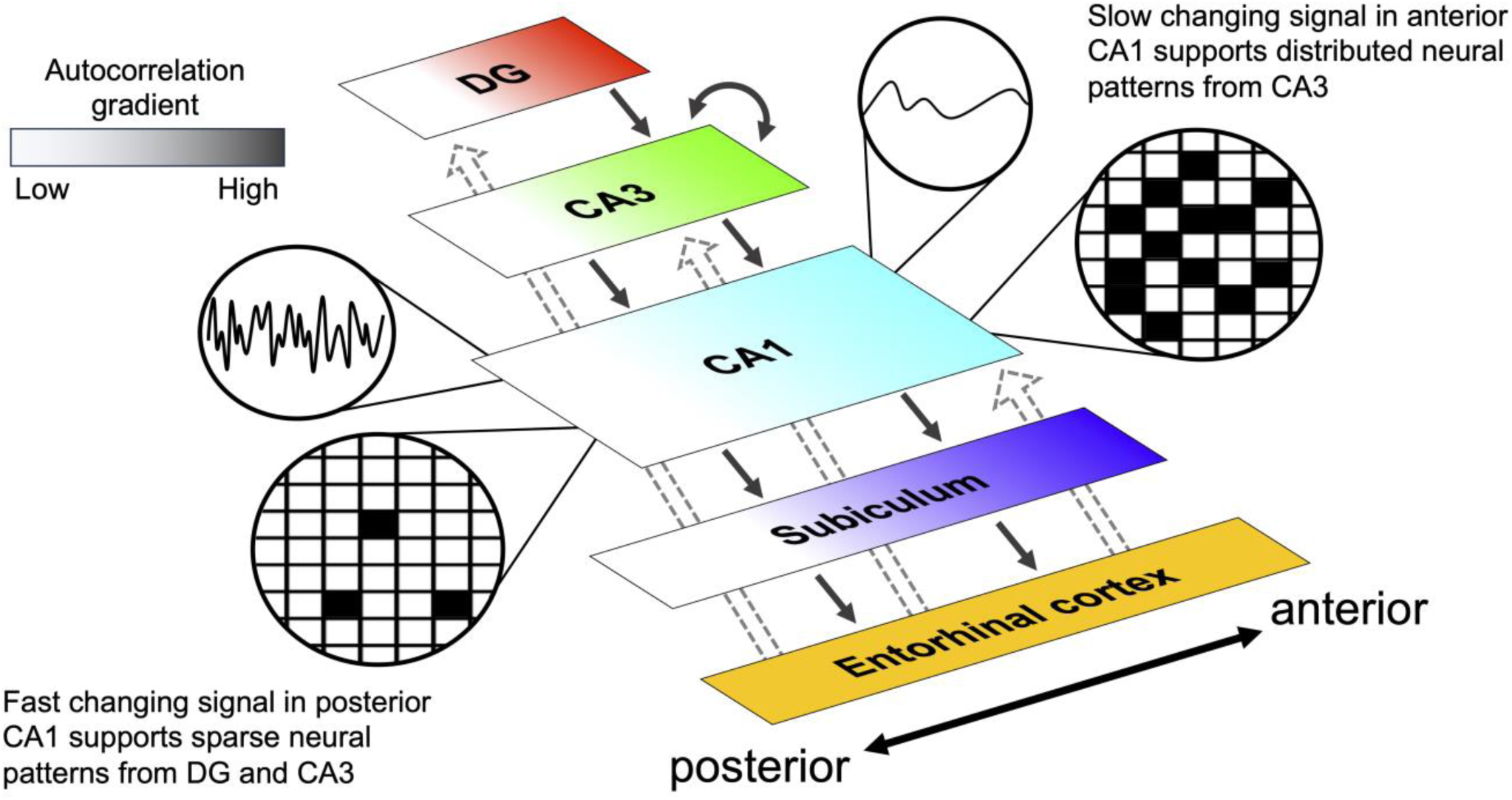
Schematic of updated hippocampal subfield model. Gradients of autocorrelation are found along the anterior-posterior axis of each subfield (represented by graded color in the figure). Slow changing signal in the anterior CA1 supports distributed neural patterns from CA3. Fast changing signal in posterior CA1 supports sparse neural patterns coming in from DG and CA3. These neural patterns are then passed to the subiculum and then out to the entorhinal cortex.

The subiculum was the region that was most distinct from the other subfields and had significantly higher autocorrelation compared to the rest of the subfields, while still demonstrating an anterior-posterior and medial-lateral gradient. It is possible that slow changing neural activity we observed in the subiculum is necessary for integrative processes that underlie holistic scene processing or scene construction typically associated with the subiculum (Dalton & Maguire, 2017; Zeidman et al., 2015). We can incorporate these subiculum findings to our new model of the hippocampal subfields and make predictions that the subiculum gradient might be supporting more integrated representations in the anterior portion (high autocorrelation) and more distinct representations in the posterior portion (low autocorrelation). Support for this hypothesis comes from a human fMRI study that parcellated the subiculum based on co-activation with the rest of the brain (Chase et al., 2015). They found a bilateral anterior subiculum cluster, where the anterior subiculum in both hemispheres was assigned to one cluster, suggesting high similarity in their co-activation profiles (Chase et al., 2015). They also found distinct left intermediate and left posterior clusters that had different coactivation from the right intermediate and right posterior (Chase et al., 2015). It is possible that integration patterns in the anterior subiculum are similar and overlapping in the left and right hemispheres, whereas neural patterns in the intermediate and posterior subiculum are more distinct and therefore differ between left and right hemispheres.

We examined the overlap between data-driven autocorrelation clusters and the anatomical subfield segments. Subfields that have higher autocorrelation should have more overlap with the anterior-medial cluster, the high autocorrelation cluster, and subfields that have lower autocorrelation should have more overlap with the posterior-lateral cluster, the low autocorrelation cluster. Indeed, we found that CA1 had lower autocorrelation and greater overlap with the posterior-lateral cluster and subiculum had higher autocorrelation and greater overlap with the anterior-medial cluster. These findings are also consistent with Plachti et al., (2019) who examined overlap between hippocampal subfields and data-driven clusters based on hippocampal resting state functional connectivity. In their study, they found that subiculum had overlap with an anterior-medial cluster, and that CA1 had overlap an intermediate lateral and posterior-lateral cluster.

We aimed to replicate the group-level autocorrelation clusters found in Bouffard, Golestani et al., (2023) and found that, in this dataset, only the right hemisphere clusters were organized into the expected anterior-medial and posterior-lateral clusters, whereas the left hemisphere clusters did not follow this pattern. In our examinations of autocorrelation gradients at the individual-level, we also replicated the anterior-posterior gradient in the right hemisphere but did not find an anterior-posterior gradient in the left hemisphere. Hemispheric differences in data-driven hippocampal clusters have been found before (Chase et al., 2015; Plachti et al., 2019; Thorp et al., 2022), so it is possible that there are inherent differences between hemispheres that we were only able to uncover using a high-resolution fMRI. Here we used a resting state dataset, as opposed to a dataset in which participants were completing a hippocampally-dependent task. It is possible that the hippocampal subfields demonstrate different autocorrelation profiles when performing tasks that recruit the hippocampus compared rest. Future research using high-resolution fMRI during a hippocampally-dependent task should be done to understand whether this hemispheric difference is a reliably characteristic of hippocampal temporal dynamics during rest and task.

Evidence from human and rodent work suggests that hippocampal subfields are not homogenous structures and that there are neural and molecular gradients that extend throughout each subfield. Prior work from human neuroimaging found anterior-posterior gradients of functional connectivity in the subiculum, CA1-3, and CA4/DG (Dalton, et al., 2019; Vos de Wael et al., 2018). Single unit recordings in the rodent hippocampus have also found distinct neural gradients within subfields, where there is a dorsal-ventral gradient of signal firing patterns in CA1 and CA3 (Barnes et al., 1990; Jung et al., 1994; Keinath et al., 2014; Kjelstrup et al., 2008; Komorowski et al., 2013; Leutgeb et al., 2004; Maurer et al., 2005). Research investigating the gene expression of cells within the rodent hippocampus also reveal distinctions along the long axis. DG and CA1 for example are segregated into three major molecular domains (dorsal, intermediate, ventral) and CA3 can be divided into 9 molecular domains (Strange et al., 2014). The results from the present study are consistent with these findings and demonstrate continuous gradients of autocorrelation along the anterior-posterior and medial-lateral axes that are recapitulated within each subfield. These findings support the notion that a close consideration of the variation of neural processes along the long axis will be necessary for full characterization of hippocampal subfield function and organization in future models.

## Conclusion

We analyzed high-resolution resting state fMRI data and found anterior-posterior and medial-lateral gradients of autocorrelation within individual hippocampal subfields. These findings add to integrative model of hippocampal organization by bridging both continuous and discrete components and challenge the notion that hippocampal subfields are homogenous units that performs a singular function. Instead, temporal dynamics vary along anterior-posterior and medial lateral axes of each subfield. This work opens the door for future research to empirically test how neural fluctuations along the long axis of hippocampal subfields are related to cognition.

